# MARSHAL, a novel tool for virtual phenotyping of maize root system hydraulic architectures

**DOI:** 10.1101/798975

**Authors:** Félicien Meunier, Adrien Heymans, Xavier Draye, Valentin Couvreur, Mathieu Javaux, Guillaume Lobet

**Affiliations:** Earth and Life Institute-Environment, UCLouvain, Belgium; Computational and Applied Vegetation Ecology lab, Ghent University, Belgium; Earth and Life Institute-Agronomy, UCLouvain, Belgium; Agrosphere (IBG3), Forschungszentrum Jülich GmbH, Germany

**Keywords:** Maize root system, root hydraulic conductivities, Root system macroscopic properties, Dynamic root system hydraulic architecture, Water flow in the soil-plant-continuum

## Abstract

Functional-structural root system models combine functional and structural root traits to represent the growth and development of root systems. In general, they are characterized by a large number of growth, architectural and functional root parameters, generating contrasted root systems evolving in a highly nonlinear environment (soil, atmosphere), which makes unclear what impact of each single root system on root system functioning actually is. On the other end of the root system modelling continuum, macroscopic root system models associate to each root system instance a set of plant-scale, easily interpretable parameters. However, as of today, it is unclear how these macroscopic parameters relate to root-scale traits and whether the upscaling of local root traits are compatible with macroscopic parameter measurements. The aim of this study was to bridge the gap between these two modelling approaches by providing a fast and reliable tool, which eventually can help performing plant virtual breeding.

We describe here the MAize Root System Hydraulic Architecture soLver (MARSHAL), a new efficient and user-friendly computational tool that couples a root architecture model (CRootBox) with fast and accurate algorithms of water flow through hydraulic architectures and plant-scale parameter calculations, and a review of architectural and hydraulic parameters of maize.

To illustrate the tool’s potential, we generated contrasted maize hydraulic architectures that we compared with architectural (root length density) and hydraulic (root system conductance) observations. Observed variability of these traits was well captured by model ensemble runs We also analyzed the multivariate sensitivity of mature root system conductance, mean depth of uptake, root system volume and convex hull to the input parameters to highlight the key parameters to vary for efficient virtual root system breeding. MARSHAL enables inverse optimisations, sensitivity analyses and virtual breeding of maize hydraulic root architecture. It is available as an R package, an RMarkdown pipeline, and a web application.

**One-sentence summary:** We developed a dynamic hydraulic-architectural model of the root system, parameterized for maize, to generate contrasted hydraulic architectures, compatible with field and lab observations and that can be further analyzed in soil-root system models for virtual breeding.

**Authors contributions:** F.M., X.D., M.J. and G.L. designed the study and defined its scope; F.M. and G.L. developed the model while associated tools were created by A.H. and G.L.; F.M. ran the model simulations and analyzed the results together with M.J and G.L.; F.M. and M.J. wrote the first version of this manuscript; all co-authors critically revised it.

## Introduction

Root systems (RS) are excellent candidates for contributing to the development of novel drought tolerant cultivars because of their key role in water uptake and their wide genetic variability (Wasson et al. 2012). Several authors, such as Lynch (2013) for maize or Comas et al. (2013) for wheat, have proposed specific combinations of optimal trait states that should minimize water limitation and potentially optimize yield under scarce water conditions (Donald 1968). However, many traits conferring drought tolerance present dual effects as a function of water availability and climatic demand time (Leitner et al. 2014; Francois Tardieu, Draye, and Javaux 2017; François Tardieu 2012). Furthermore, in many cases, impacts of specific traits are studied alone and it is not always clear where combinations of potentially positive traits would lead to in actual field conditions.

To clarify the picture, functional-structural root system models (FSRSM) have a crucial role to play. Today, state-of-the-art FSRSM are able to numerically simulate water movement in the so-called soil-plant-atmosphere continuum from the root uptake location to the shoot. These models combine root functional (i.e. hydraulic) and structural (i.e. architectural) information to describe plant root systems and represent their functioning over time in their surrounding environment (soil and atmosphere), see for example R-SWMS from Javaux et al. (2008) or OpenSimRoot from Postma et al. (2017).

FSRSM are potentially useful instruments for virtual breeding as they allow one to investigate the behavior of any desired combination of plant traits within any environmental condition, including contrasted pedo-climatic scenarios (Chapman 2007). They also allow one to highlight mechanisms, regulation rules and feedbacks that are expected to influence plant transpiration (Huber et al. 2015) and yield (Meunier et al. 2016; Srinivasan, Kumar, and Long 2016). In addition, FSRSM enable crossing scales and to predict the impact of deterministic relations, observed and valid on short timescales, on the overall relations within a system in specific pedo-climatic conditions (Hammer et al. 2009).

In theory, thus, it is possible to virtually generate root system ideotypes for water uptake by running thousands of instance of a FSRSM over successive crop seasons (Jubery et al. 2019). In practice, unfortunately, the heavy computational load, the high time requirement for running a single simulation in 3D explicit numerical soil-plant models, and the huge problem dimensionality (i.e. the number of model parameters to vary) do not allow such models to be routinely used to predict plant performance in contrasted pedo-climatic conditions yet.

Fast and simple approaches have been used recently to solve root water uptake equations (Javaux et al. 2013). They rely on upscaled models (Couvreur et al. 2012; de Jong van Lier et al. 2008), which do not describe the RSHA and surrounding soil matrix as a complex ensemble of local properties. Instead, they represent them as a reduced number of macroscopic root properties (i.e. K_rs_ the root system conductance, and SUF the uptake profile in homogeneous conditions) that determine water flow rates at the soil-root interfaces with satisfactory accuracy and decreased simulation times operational for land surface models (Sulis et al. 2019). These macroscopic RSHA properties can be derived from local root traits using ad-hoc plant hydraulics algorithm (Meunier et al. 2017), by inverse modelling of soil moisture dynamics data (Cai et al. 2018), or can even be measured in the lab. However, as of today, it is not clear if the upscaling of local root traits are compatible with RS-scale parameter measurements. In addition, macroscopic parameter sensitivity to root local traits was never tested because of the lack of appropriate tools.

A model to investigate the combined impact of plant development and hydraulic root traits on effective plant scale hydraulic properties is thus missing. In this study, we present MARSHAL, the MAize Root System Hydraulic Architecture soLver that combines the root architecture model CRootBox (Schnepf et al. 2018) with the hybrid solution for water flow in RSHA of Meunier et al. (2017) to compute the macroscopic parameters of Couvreur et al. (2012). MARSHAL calculates root system conductance, instantaneous 1 to 3-D water uptake distribution and other upscaled variables (plant leaf water potential, root system volume, convex hull volume etc.) for any combination of structural and functional traits under heterogeneous and static environmental boundary conditions. In addition, MARSHAL can also estimate the evolution of plant scale properties with plant ageing thanks to the age-dependency of hydraulic properties and its root growth module.

In order to evaluate the performance of the model, we developed two case studies. In the first one, we used MARSHAL to generate an envelope of potential trajectories of maize root length densities and root system conductances with age. These envelopes were compared to observations from the literature as part of a broader meta-analysis to demonstrate the model plausibility. In the second case study, we investigated how sensitive plant-scale parameters are to local root hydraulic, growth and architectural parameters and, by doing so, we showed how the optimization of RSHA in a coupled soil-plant model could be efficiently eased.

Maize was chosen as illustrative crop species for MARSHAL because of the abundant literature on its architectural and hydraulic properties (over time). However, the workflow itself (literature meta-analysis, model calibration, runs and validation) is generic and could be applied to any plant species provided that data on growth, hydraulic and architectural traits is available. In the past, growth and architectural have been typically obtained using literature data (Kutschera 2010) or experimental observations in rhizotrons (Claude Doussan et al. 2006), minirhizotrons (Cai et al. 2016), 3D tomography (Flavel et al. 2017) or shovelomics (Trachsel et al. 2010; Zhu et al. 2011). Model required hydraulic properties can be either characterized by direct measurements (Bramley et al. 2009; Knipfer and Fricke 2011) or by combining experiments and FSRSM in an inverse modeling scheme (Doussan 1998; Meunier et al. 2017; Meunier et al. 2018; Zarebanadkouki et al. 2016; Passot et al. 2019).

### Description of MARSHAL

MARSHAL is the combination of the root architecture model CRootBox (Andrea Schnepf et al. 2018), the model of water flow in RSHA of Meunier et al. (2017), and the model of Couvreur et al. (2012). These three building blocks are summarized in the respective sections below.

#### Maize root architectural model and parameters

CRootBox considers several root types. A set of parameters is associated to each root type and defined as a combination of a mean and a standard deviation. For each root, basal, branching, and apical zones are distinguished. New lateral roots are only created when the basal and apical zones have developed their full length. Then each lateral emerges at a regular interbranching distance d_inter_, with a maximal number of lateral roots N_ob_ with an insertion angle θ between the parent root and the new lateral root (Figure 1). Tropisms are also implemented in the model under the form of change of heading angles. Each root type has a constant radius r.

**Figure 1:**
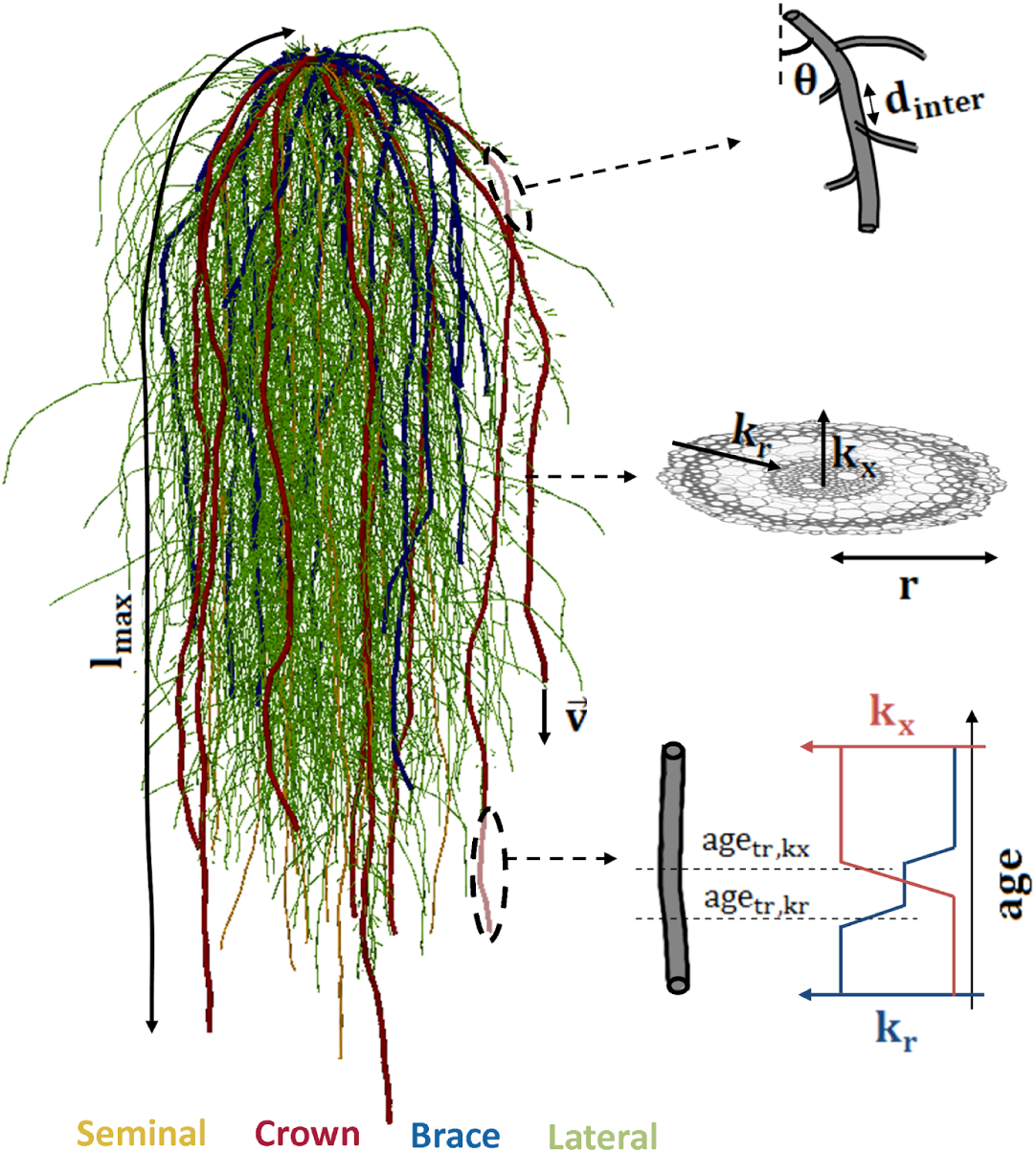
Maize root system architecture as generated by MARSHAL. Each root type (i.e. seminal, crown, brace and each type of lateral root) is characterized by a set of architectural and hydraulic parameters defined in Table 1. Root system-scale hydraulic and geometrical parameters can be calculated from each model instance (Table 1).

The nomenclature of maize root types follows Hauck et al. (2015). The development of the root system is divided into two stages (embryonic and post-embryonic growth), as observed by Hochholdinger et al. (2004). The former is constituted by the root primary and seminal roots emerging from the seed alongside with their laterals. The post-embryonic root system starts developing around one week after the emergence of the primary and seminal roots and becomes prominent approximately 2 weeks after seed germination. It is made of crown roots, i.e. shoot-borne roots that develop from nodes below the soil surface and brace roots, shoot-borne roots developing from nodes above the soil surface several weeks later as the plant matures. These two types of roots also exhibit lateral branching, although the brace roots do not naturally branch until they penetrate the soil.

**Table 1:** Root, root system and root system macroscopic parameters. L = Length, P = Pressure, T = Time. Model actual models units are cm (L), hPa (P) and day (T).

| Name | Dimension | Description |
| --- | --- | --- |
| **Root type parameters** |  |  |
| v | L T-1 | Initial elongation rate |
| lmax | L | Maximal single root length |
| r | L | Root radius |
| $\theta$ | - | Insertion angle |
| dinter | L | Inter-branching distance |
| Nob | - | Maximal number of lateral roots |
| kr | L T-1P-1 | Root radial conductivity |
| kx | L4T-1P-1 | Root axial conductance |
| akx | T | Transition ages for root axial conductance |
| agkr | T | Transition ages for root radial conductivity |
| **Root system parameters** |  |  |
| Nmax | - | Maximal number of main axes |
| t | T | Starting time of crown or brace root development |
| z | L | Vertical position of crown and brace root development |
| **Root system macroscopic parameters** |  |  |
| VCH | L3 | Convex hull volume of the root system |
| VRS | L3 | Root system volume |
| Krs | L3T-1P-1 | Root system conductance |
| zSUF | L | Mean depth of root water uptake |

The main root axes of the maize plant are implemented as crown or brace roots (in addition to the primary and seminal roots) that start to grow after predefined times and predefined depths (z_I_ vertical position on the stem). There is a maximal total number of main root axes N_max_.

In total, five root types are considered for maize root systems in MARSHAL: primary and seminal (considered together), short and long laterals, crown and brace roots. Lynch (2013) reviewed ranges of observations of most of the architectural parameters defined above and that could be used as input variability ranges in the model. Table 1 summarizes the root-type parameters and the root system parameters, respectively.

#### Maize root hydraulic model and parameters

The water flow model included in MARSHAL is the water flow algorithm developed by (Doussan 1998), improved by including the analytical solution of Landsberg and Fowkes (1978) extended for root systems by Meunier et al. (2017). The model solves the water flow equations in hydraulic architecture under user-prescribed boundary conditions (stem water potential Ψ _*collar*_) and water potentials at all soil-root interfaces (Ψ_*sr*_). Numerically, the solution provided to the user is the xylem water potential Ψ_*x*_ at the basal end of each root segment (from 1 to N_seg_), each segment being characterized by a root radial hydraulic conductivity (k_r_ for the water pathway from soil-root interface to xylem vessels) and a specific axial hydraulic conductance (k_x_ for the axial water pathway in xylem). These properties may vary with root type and segment age at prescribed transition ages (age_tr_). The list of hydraulic parameters can also be found in Table 1 and are further illustrated in Figure 1. When the RSHA is determined, the water flow problem is solved by calculating:

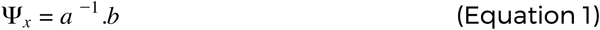

where a [matrix of dimension N_seg_ × N_seg_] and b [vector of dimension N_seg_ × 1] are built based on the root system architectures generated over time by CRootBox and the root properties (see Meunier et al. 2017 for more details). b contains the top-boundary condition, i.e. the plant collar potential Ψ_*collar*_ and the soil-root interface water potentials (multiplied by a hydraulic conductance) while a contains the segment connections and their hydraulic properties. Based on each segment’s Ψ_*x*_, MARSHAL calculates the water flow within the root system. In particular, water radial flow in each segment (J_r_ ) is given by:

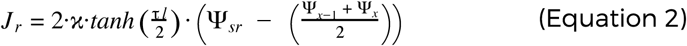

where *κ* (defined as 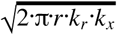) and *τ* (defined as 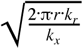 are hydraulic-specific root segment properties, see Meunier et al. (2017) and Ψ_*x*−1_ is the water potential at the proximal end of each segment (Equation 1). Note that MARSHAL is insensitive to root segment size (provided that their properties are uniform within segment), since it uses an analytical solution of water flow within infinitesimal sub-segments (Meunier et al. 2017).

Several studies measured hydraulic axial and radial conductivities for different maize root types at different ages and for different genotypes. Meunier et al. (2018) performed a literature meta-analysis and defined the genetic range of variation of axial and radial hydraulic per root type, which could also be used in the model as input variability ranges.

#### Root system macroscopic parameters

After generating the maize RSHA according to the user input parameters, MARSHAL calculates a set of macroscopic parameters, in particular those of Couvreur et al. (2012). Those are the root system conductance K_rs_ (from soil-root interfaces to xylem vessels at the plant collar) and the Standard Uptake Fractions SUF (i.e. the relative contribution of each root segment to the total water uptake under homogeneous soil water potential conditions). Solving Equations 1-2 under uniform Ψ_*sr*_ and with a pre-definite collar potential Ψ_*collar*_ provides radial inflow J_r_, which allows MARSHAL to calculate these parameters as:

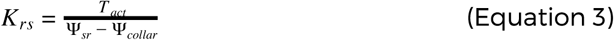

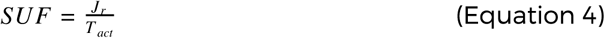

where T_act_ is the sum of water flow rates over all root segments i:

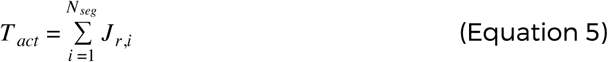

SUF is then further used to generate the mean depth of uptake as the weighted mean of the root water uptake depths (Meunier et al. 2016):

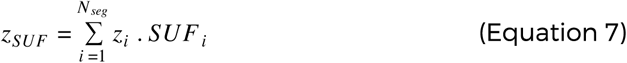

where *z*_*i*_ is the i^th^ root segment depth.

In addition, for each model run, MARSHAL calculates several root system scale geometrical properties including the root system convex hull volume (V_CH_, i.e. volume of the smallest convex set that entirely contains the root system), the root length density profile (RLD, i.e. root length per unit soil volume), and the total volume of the root system (V_rs_, as the sum of individual root segment volumes).

#### MARSHAL is freely available and highly flexible

MARSHAL is available as an R package (doi: 10.5281/zenodo.2391555). It was embedded into both an RMarkdown pipeline (for batch analysis, publicly available at https://github.com/MARSHAL-ROOT/marshal-pipeline, doi: 10.5281/zenodo.2474420) and a Shiny web application (https://plantmodelling.shinyapps.io/marshal/, 10.5281/zenodo.2391249). Figure 2 illustrates the overall model workflow. Reference maize root system hydraulic architectures are generated using literature hydraulic and architectural parameters. The default growth and architectural parameterization of the online version of MARSHAL is a set of parameters for that could reproduce observed dynamics of maize root system architectures (Ahmed et al. 2018). Root hydraulic properties used as default are the ones obtained by Doussan et al. (1998), see in particular Figure 4 in the aforementioned paper. Each single model parameter can be modified by the user, which results in hydraulic architectures visible almost instantaneously.

**Figure 2:**
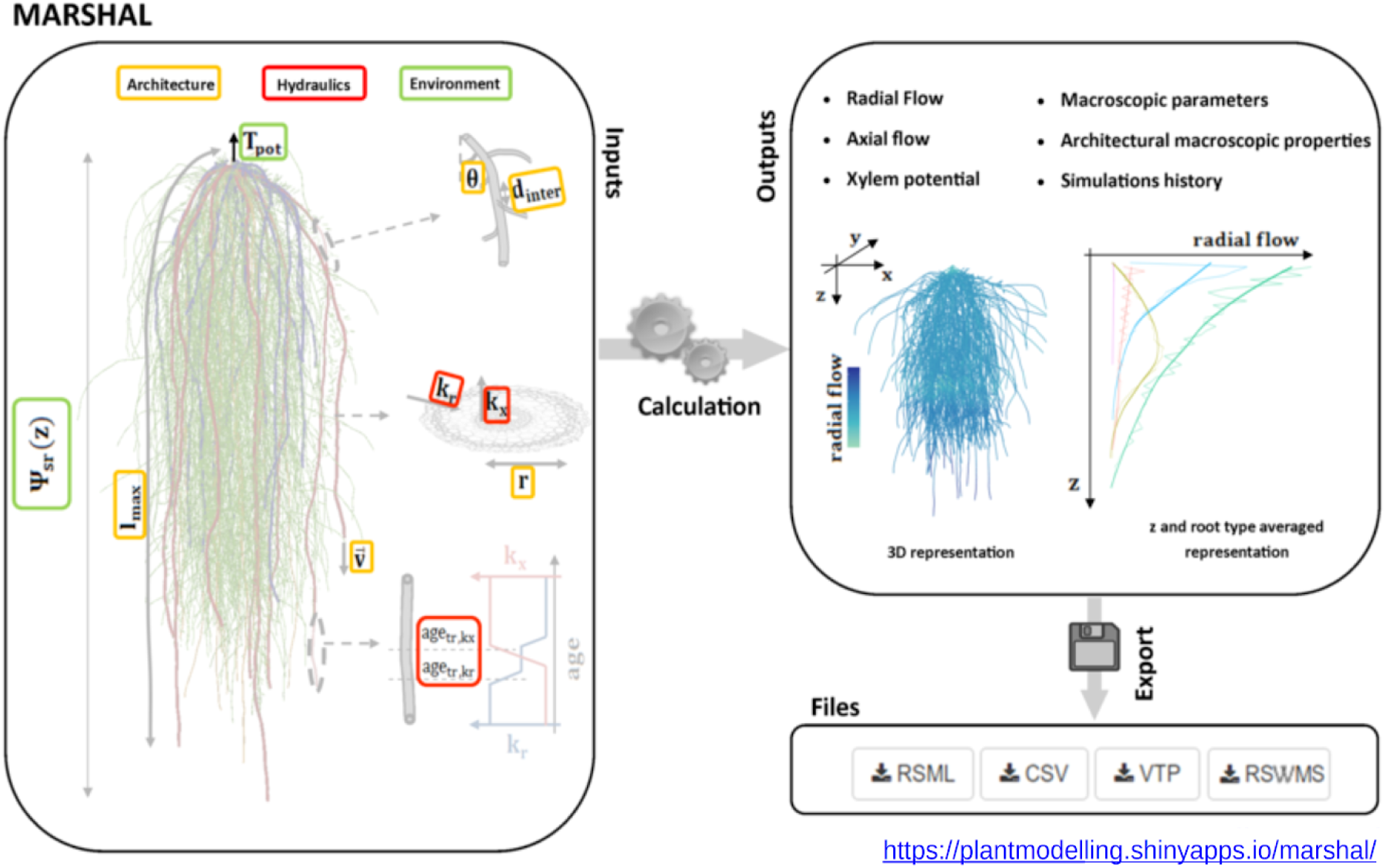
Schematics of MARSHAL workflow. First, model input parameters (related to the RS architecture and hydraulics, as well as the environment) are provided by the user. MARSHAL then generates the hydraulic architecture and computes the water flow and the macroscopic properties. The generated RSHA can be visualized in 3D or with the help of depth-averaged plots. Eventually, root system metrics and RSHA can be exported using standard formats for further use (among other in soil-root models)

Hydraulic, growth and architectural parameter modification affects the water uptake location and water flow within the root system, which may also be seen from 3D plot of the root system or z-averaged profiles of water flow (Figure 2, right panel). Finally, the profile of soil-root water potential may also be changed to assess the impact of soil water potential heterogeneity on root water uptake distribution. In any case, Equations 1-2 are always solved under both prescribed and homogeneous soil water potential conditions to calculate root system macroscopic properties (Equations 3-6).

All root system metrics such as root length density, or total root length and surface are made available to the user for export. The online version of MARSHAL also saves previous simulations outputs during the same session, allowing users to easily compare the effect of changes in input parameters. Finally, when a root system hydraulic architecture is suitable for the user, it can be easily exported in a common format, e.g. RSML file for example (Lobet et al. 2015). Root system macroscopic properties can also be exported for further use in soil-plant coupled model.

## Material and Methods

Two case studies illustrate some of the model potentialities. The first one compares literature data with ranges of root system scale architectural and hydraulic properties predicted from local root traits. The second one analyses the sensitivity of root system scale properties to local root traits.

### Comparison of model predictions with experimental data

Ten thousand maize root system architectures were first generated in a parametric space defined as MARSHAL default parameter set (Table 2) and a coefficient of variation of 50% for each single parameter. The panels of root systems was then used to compare model predictions of root system conductances and root distributions with actual data.

**Table 2:** Model default architectural parameters (see Table 1 for description and units and Schnepf et al. (2018) for details).

| **Root type parameters** |  |  |  |  |  |
| --- | --- | --- | --- | --- | --- |
| **Param** | **Primary/seminal** | **Crown** | **Brace** | **Short lateral** | **Long laterals** |
| la | 15 | 15 | 13 | 1 | 3 |
| lb | 2 | 2 | 2 | 1 | 2 |
| ln | 1 | 1 | 0.5 | 1 | 1 |
| nob | 96 | 63 | 270 | 1.5 | 30 |
| v | 1.1 | 1.6 | 1 | 1.3 | 1.13 |
| r | 0.12 | 0.1 | 0.2 | 0.04 | 0.04 |
| θ | 0.97 | 1.1 | 1.1 | 1.13 | 1.13 |
| successor | 5 | 4 | 4 | - | - |
| **Root system parameters** |  |  |  |  |  |
| Nmax | 10 (crown), 18 (brace) |  |  |  |  |
| t | 7 (first crown, then every 2 days), (20 first brace) |  |  |  |  |
| z | 0 |  |  |  |  |

The range of simulated root system conductances (K_rs_) with plant age was compared to ranges of measured data from literature. Maize conductance data were collected through an extensive literature search using Web of Science and Google scholar as search engines with a combination of “maize” and “root system conductance/conductivity” as keywords. Foreach single study used in this research, we collected the raw data from tables, figures or supplementary information.

The Root Mean Square of error (RMSE) between the RLD of each simulated maize root system (RLD_obs_) and independent field observations (RLD_obs_) by Gao et al. (2010) was then computed. The comparison was carried out at different times t and horizontal distances to the stem (0, 10, and 20 cm, perpendicularly to the row). To account for the impact of neighboring plants on RLD_obs_, copies of each root system were located according to seed positions in the field (distant of 30 and 50 cm in the direction parallel and perpendicular to the row, respectively). RMSE was calculated as:

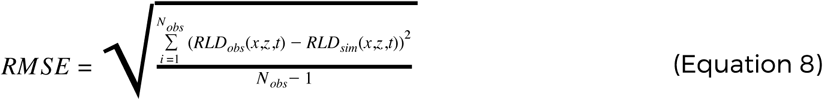

with N_obs_ the total number of observations in time (time and in the two-dimensional space x-z). The generated root systems were then ranked according to their RMSE. RS hydraulic parameters were then re-calculated using the default set of hydraulic parameters (and no variation) to evaluate the impact of architectural parameters only on plant-scale hydraulic parameters.

### Model parameter sensitivity

A set of 100 000 root systems was then generated with MARSHAL to test the impact of local root trait changes on root system scale properties based on the parametric space defined above (default ± 50%). The global sensitivity algorithm of Cannavó (2012) was used to compute the sensitivity of macroscopic properties to local root traits. The analyzed plant-scale properties were the root system conductance (K_rs_), the depth of standard uptake (z_SUF_), the root system volume (V_rs_) and the convex hull volume (V_CH_), calculated 60 days after sowing.

The explanatory local root traits were gathered by parameter family (either hydraulic or architectural). They were then assembled according to their corresponding root types (brace, lateral, etc.) or parameter type (insertion angle, elongation rate, etc.). The measured sensitivity in this analysis is the contribution to the output variance explained by the parameter family, the root type or the parameter itself, calculated as the sum of the first order sensitivity index as defined by (Sobol′ 2001).

## Results and discussion

### Illustration of MARSHAL potential for root phenotyping

Figure 3 first illustrates how macroscopic properties of the virtual panel of 10 000 RSHA with parameter sets varying around default values (coefficient of variation of 50%) evolve in time through growing and ageing. Relationships between root system conductance K_rs_ and root system volume V_rs_ (Figure 3a), depth of uptake z_SUF_ and conductance K_rs_ (Figure 3b) and convex hull volume V_CH_ and root system V_rs_ volumes (Figure 3c) are displayed for all simulated root systems (scatter plot) at all times (colours). In addition, the relationship between average values with root system aging is shown as a solid dark line. Probability density functions of macroscopic properties are also displayed alongside panel axes with the same color code (root systems age).

**Figure 3:**
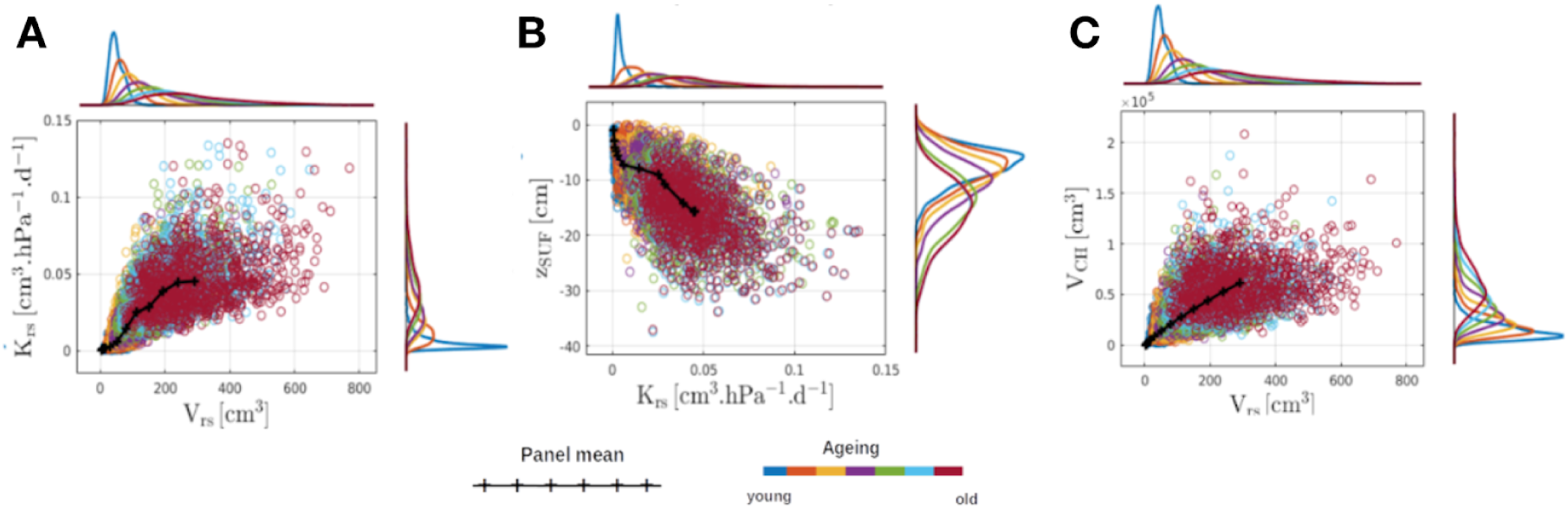
Example of maize virtual panel generation (N = 10 000). The three panels represent the relationships between root system hydraulic and/or architectural properties for a large number of maize individuals as they age (see Table 1). The resulting density distributions are also represented over time with the same colour scale.

The initial narrow distributions expand with root system aging since the first model outputs are by definition well constrained (they all start from a seed with no roots). Model input variability translates into widespread values of macroscopic properties. As these properties partly determine the tolerance to water deficit (Alsina et al., 2011; Schoppach et al., 2014), they could be optimized (Meunier et al. 2016, Leitner et al. 2014), thereby decreasing the dimensionality of the optimal RSHA problem (i.e. one could optimize the root system conductance K_rs_ and other root macroscopic properties rather than all particular local conductivities and the full root system map).

### MARSHAL predicts reliable plant root system conductance and architecture as compared to experimental data

Figure 4 compares the range of simulated K_rs_ when exploring the dynamic architectural and hydraulic parameter spaces (blue enveloppe) with measured root system conductances from independent experiments (grey boxes are experiment data from (Zhang, Zhang, and Liang 1995; Zhuang et al. 2001; Smith and Roberts 2003; Parent et al. 2009; Hu et al. 2011; Wang et al. 2013; Caldeira et al. 2014; Meunier et al. 2016)).

**Figure 4:**
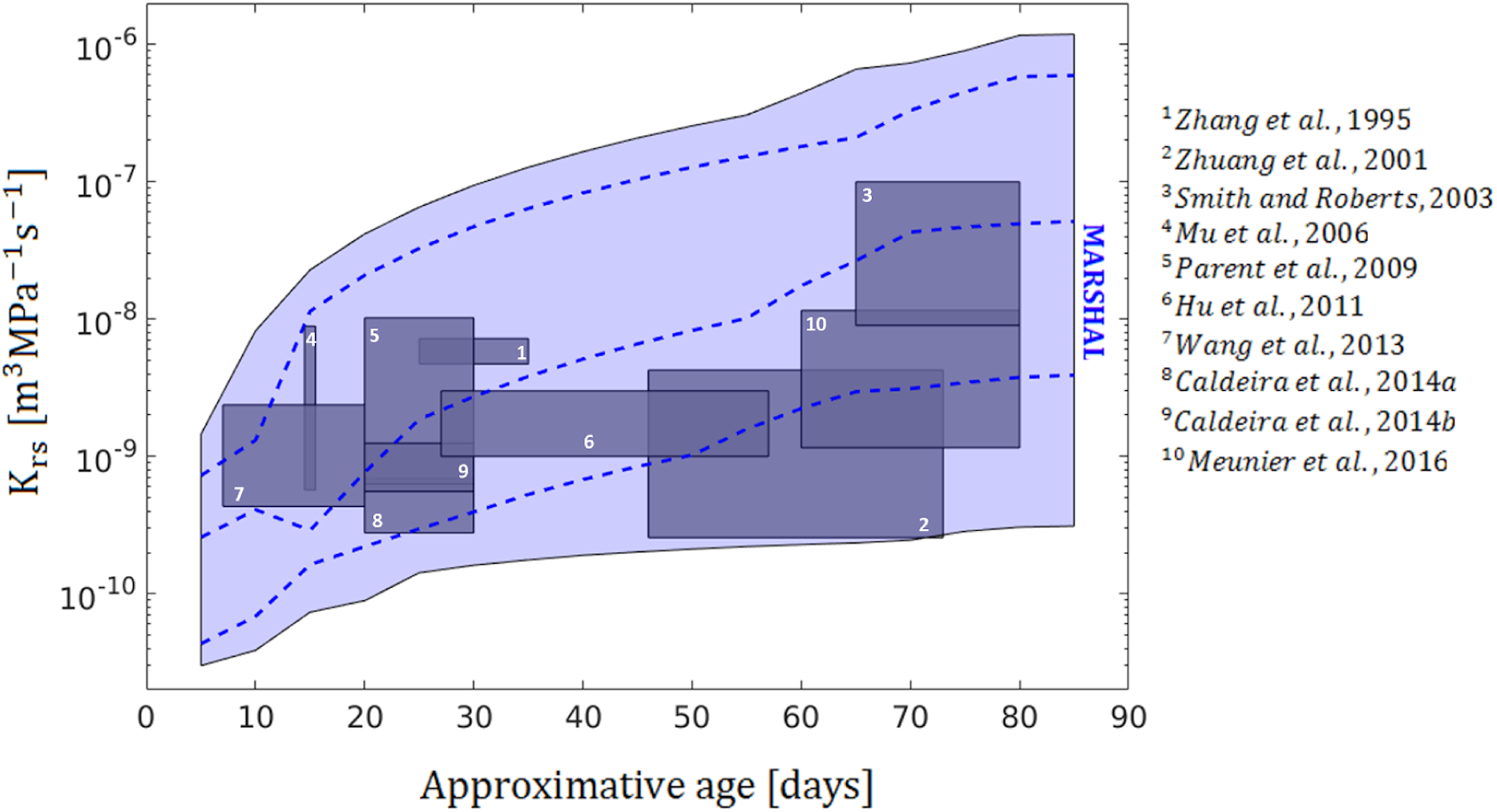
Comparison between maize root system conductances observed in the literature (grey boxes) and the range of conductances simulated using MARSHAL (blue envelope) over the full crop cycle. Dashed lines are three particular realizations of root system conductance trajectories.

Observational data traits are presented as boxes as different root system ages and treatments in these studies result in variability in the x and y axes, respectively. Interestingly, orders of magnitude between observed and simulated conductances are similar and all observations (in grey) are contained in the blue envelope, suggesting that MARSHAL can bridge the gap between local root traits and plant-scale properties. The local root trait state variability is able to explain the observed variability of root system conductance.

From the same panel of 10 000 virtual root systems (phenotypes), simulated RLD were used for comparison with ones from an independent study from Gao et al. (2010). From the 10 000 estimates, we kept the 8 root systems with minimum RSME (Equation 7), see fit in Figure 5a. R-square of measured vs simulated RLD profiles were all larger than 0.9 for the 8 best simulations. For these 10 000 root systems, we further recalculated root system conductance using default hydraulic values.Interestingly, the range of variation of the best simulated root system macroscopic properties was dramatically reduced as compared to the entire variability of the panel (Figure 5b): the grey enveloppe is the whole panel range of variation of K_rs_ when only the architectural parameters change while the blue one is the range of variation of the 8 best estimates.

**Figure 5:**
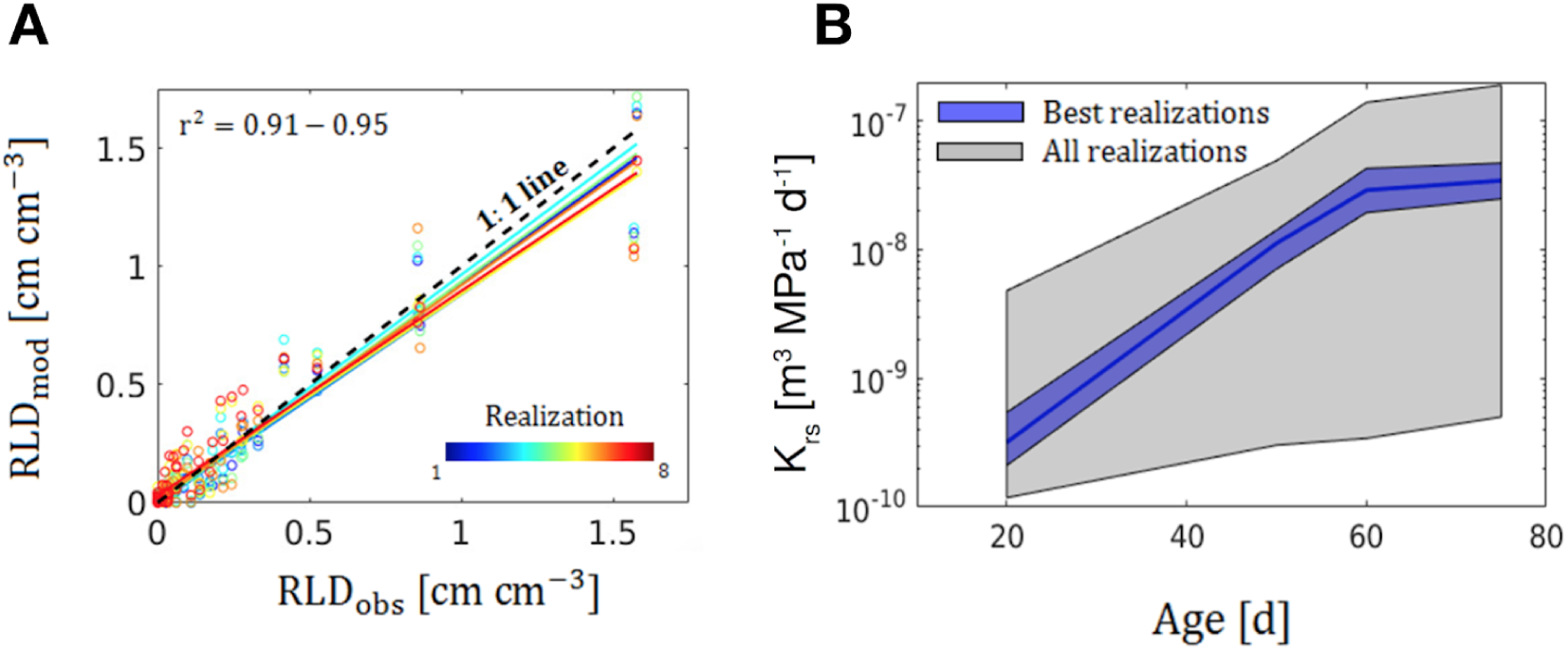
Maize root length density profile goodness of fit using MARSHAL for the 8 best realizations (A) and corresponding RS conductance change envelopes with age for the whole panel of 10 000 simulation runs (grey) and the height best simulations (blue) (B).

The problem to find a root system architecture fitting root length density profile(s) is not unique: different ensemble of trait states could lead to similar goodness of fit. However, Figure 5b suggests that these solutions could have similar values for macroscopic properties since their architecture is constrained to certain limits. This also suggests that if no unique solution is reachable in the parametric space, functional parameters could be conserved between best solutions.

Figure 5 indicates that under constraints (measurements of root length density profiles, maize root growth rules, limited parametric space), the range of variability of plant-scale properties is considerably reduced, leading to a good knowledge of the root system functioning (i.e. here the conductance). The initial variability (grey envelope) was due to the uncertainty on the total length of the root system and this information provided as the evolution of the root length densities constrained the inverse problem. The variability of plant scale properties could be reduced even further if additional metrics are used to constrain the inversion step. For instance, data obtained using the shovelomics phenotyping technique (Colombi et al. 2015; Trachsel et al. 2010), such as the number of nodal roots, could be very informative on this prospect.

In numerous studies, a gap exists between the measured metrics (conductance or root length density) and the FSRSM scales (see e.g. Trachsel et al. 2013). MARSHAL allows bridging this gap by estimating the value of observable macroscopic properties based on local root trait states. It is also able of determining parameter trade-offs and parameter uncertainties (see next section).

### MARSHAL enables sensitivity and optimization algorithms

The running time for MARSHAL is typically about 1 second on a personal laptop (tested on a processor Intel core i7 with 8Go of memory). This time is necessary to (i) generate a mature RSA (∼3-4 months) based on the actual root-type and root system parameter set, (ii) distribute the hydraulic root properties according to the root segment ages and the user-defined hydraulic properties, (iii) calculate the water flow and potential within the root system (i.e. solving Equations 1-2) and (iv) compute the plant-scale hydraulic properties. Such a short time allows one to run thousands of model instances to analyze model sensitivity to input parameters, generate run ensemble, assess the spread of plant-scale macroscopic model or optimize parameters against any objective function.

Taking advantage of the coupling between MARSHAL and global statistical algorithms to perform sensitivity analyses of the macroscopic properties, we assessed how local root traits affect (i) the root system conductance K_rs_ (Figure 6), the depth of standard water uptake z_SUF_ (Figure 7), the root system volume V_rs_ (Figure 8 left) and the convex hull volume V_CH_ (Figure 8 right) after 60 days of growth and development, corresponding on average to the flowering stage for maize. Let us note that together all parameters do not explain the entire macroscopic variance because only primary effects are plotted in the figures. Impact of combined local root traits (i.e. parameter covariances) are not accounted for in the variance decomposition.

**Figure 6:**
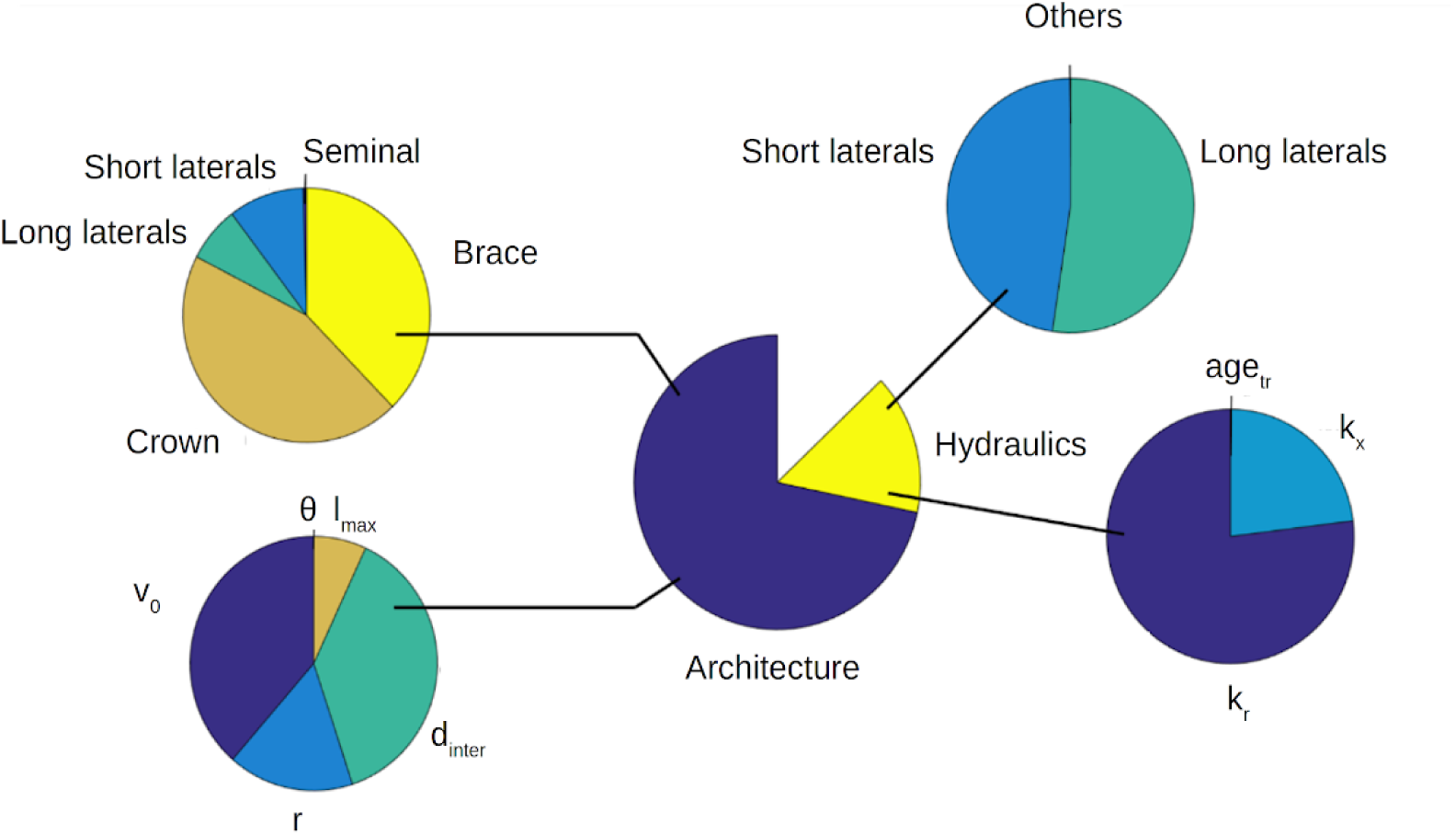
Sensitivity analysis of maize root system conductance at 60 days. The parameters are split into hydraulic and architectural parameters and classified according to the root and parameter types. For the details about parameter and root type meaning, see Figure 1 and Table 1.

**Figure 7:**
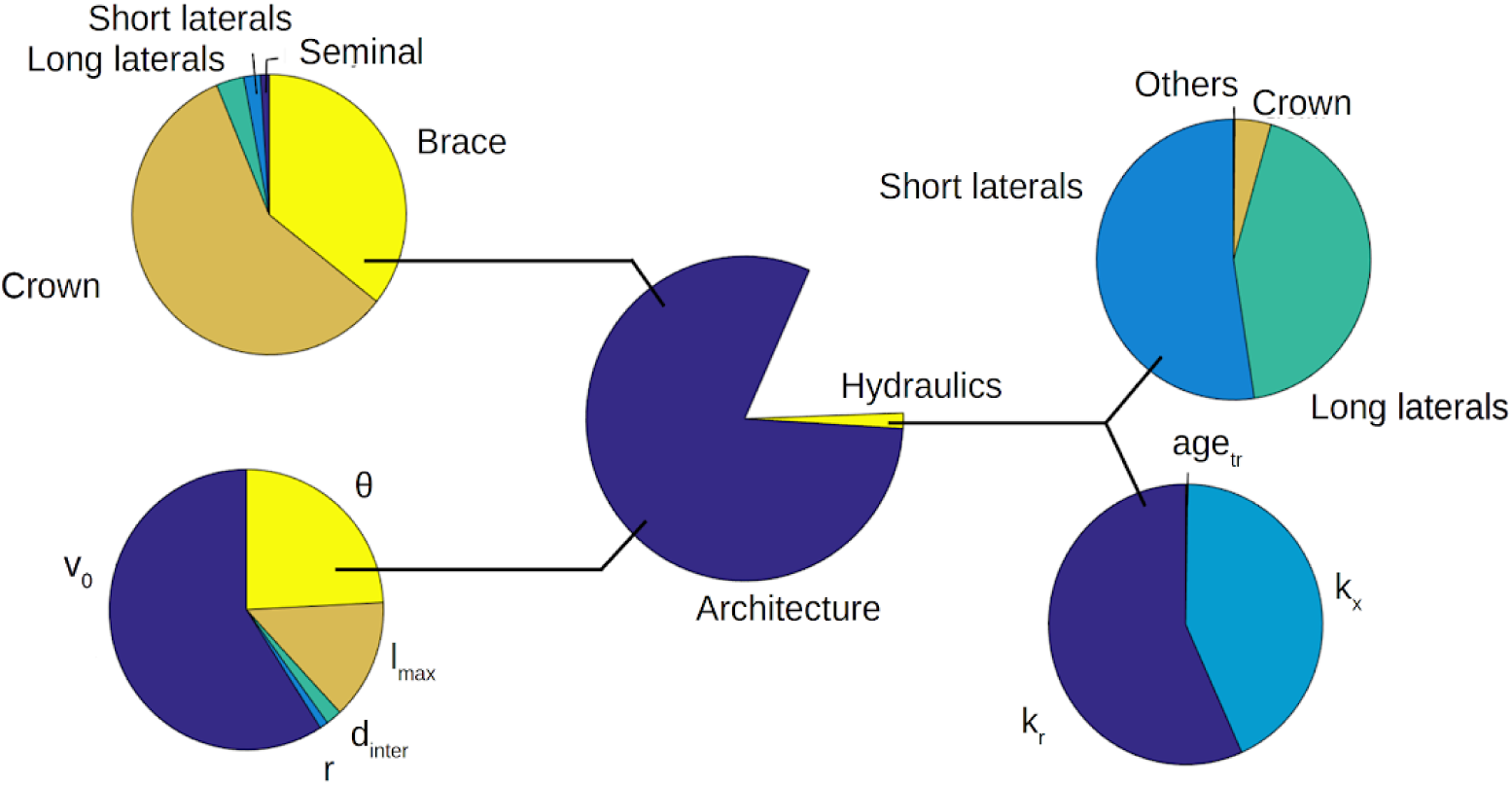
Sensitivity analysis of maize root water uptake depth at 60 days. The parameters are split into hydraulic and architectural parameters and classified according to the root and parameter types. For the details about parameter and root type meaning, see Figure 1 and Table 1.

**Figure 8:**
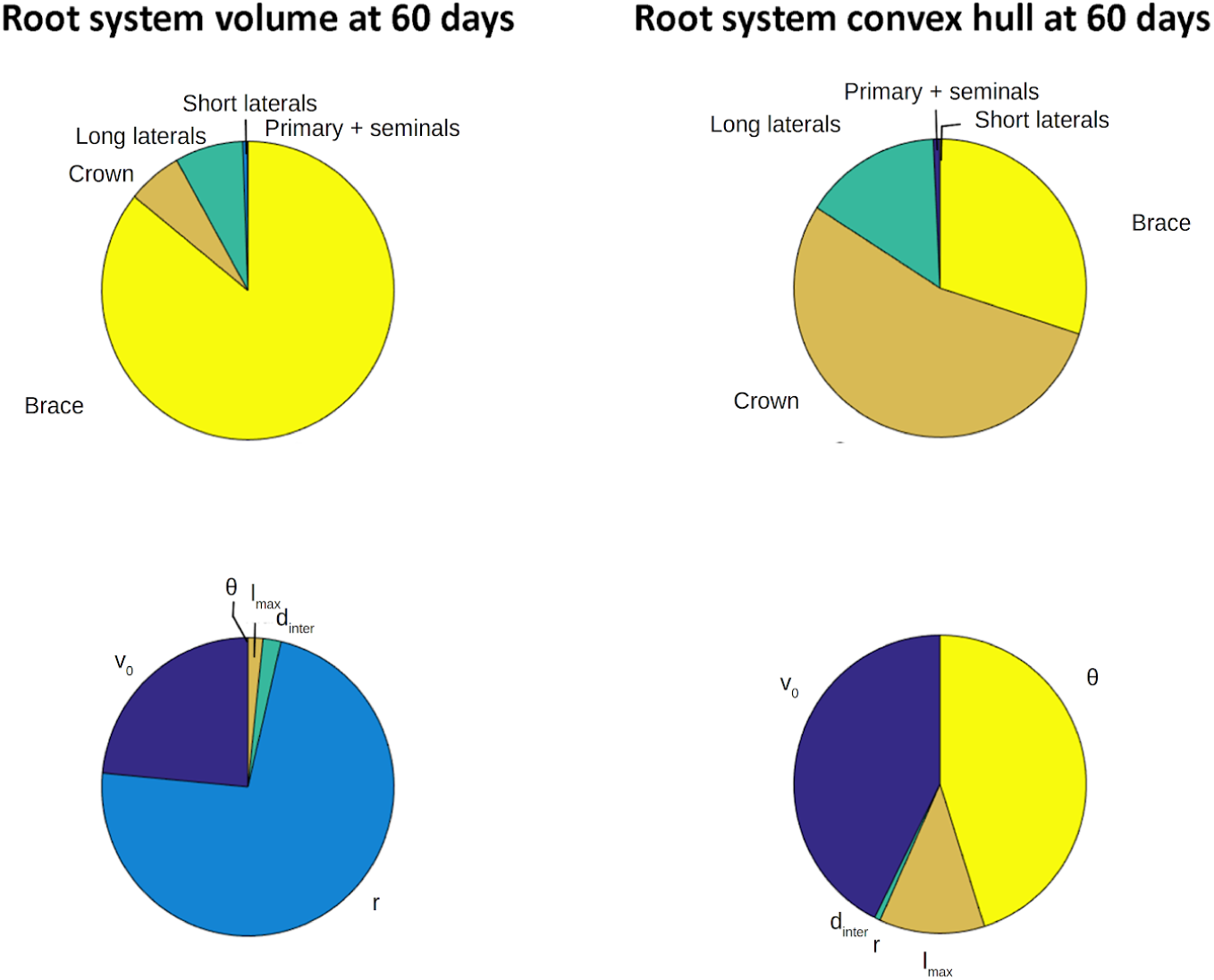
Sensitivity analysis of maize root system volume (left) and convex hull volume (right) at 60 days. The parameters are sorted according to the root and parameter types. For the details about parameter and root type meaning, see Figure 1 and Table 1.

These sensitivity analyses highlight the (in)sensitive parameters for each considered macroscopic property. For instance, insertion angles do not affect K_rs_ as only the topology and the hydraulics (not the orientation of roots) drive the value of K_rs_ (Figure 6). Yet, insertion angles of adventive roots (brace and crown) are critical for the mean depth of uptake z_SUF_ (Figure 7) as they position a large number of lateral roots that are the main facilitators of water uptake (see also Ahmed et al. 2015; Meunier et al. 2018). Hydraulics explain about 20% of the total variance of the root system conductance, which may seems low but one has to keep in mind that we used the reference value ± 50% for all parameter limits. Indeed, this is a sensitivity analysis of the model parameters, not of the potential impact of actual variation range of root traits on plant macroscopic hydraulic properties. These ranges represent much larger changes for architectural parameters than hydraulic ones as the latter have been shown to vary by several orders of magnitudes (Meunier et al. 2018). Their non-negligible contribution with such small ranges, as compared to realistic ranges, is rather an indication of their great potential to adjust plant scale hydraulic properties.

Amongst the most critical parameters affecting root system conductance, we find the elongation rates and the interbranching distances, especially of the primary roots (brace and crown) as small changes of their values would lead to dramatic changes of lateral root numbers. And as it appears that lateral roots are the main location of uptake (see Figure 6, as well as Ahmed et al. (2015)), changing their total number severely impacts the total root system conductance. The root radius (r) also plays a substantial role, in part because radial hydraulic conductances are assumed proportional to r, which is a fairly reasonable assumption (Heymans et al. 2019). As expected (Steudle et al. 1998), the main hydraulic limitation for water uptake is the radial conductivity (which explains ¾ of the variance generated by the hydraulics). Contrastingly, the axial conductances are relatively more important for the z_SUF_ even though the hydraulics as a whole does not contribute much to such metric. The insertion angles and elongation rates of primary, on the opposite, determine how deep the water is taken up by the root system in homogenous soil water potential conditions. Overall, macroscopic hydraulic properties are thus approximately as sensitive to the growth rate parameter (v0) as to all other static architectural traits together.

The architectural parameters are by definition the only one to play a role for both the volumes of the root system and the convex hull: hydraulics is actually not taken into account when computing these metrics. Interestingly, the sensitivity analyses reveal very contrasted results for these two architectural metrics. Brace roots are the main root type explanatory variable for the root system volume because of the size of their radius, which is much larger than in other root types. As root radius does not play an important role for other macroscopic properties (RS conductance for example, see Figure 6), it makes them excellent candidates for more efficient root system hydraulic architectures (similar RS conductance with a lower root system volume for example). Contrastingly, the insertion angle of brace and crown roots clearly appeared to control soil exploration (V_CH_).

### Limitations of MARSHAL

When using MARSHAL, one must keep in mind the limitations of the model, in order to properly use it and interpret its outputs accurately. The first main limitation, as stated above, is that MARSHAL does not simulate water dynamics in the soil, as would R-SWMS (Javaux et al. 2008). That said, any heterogeneous soil water potential profile can be used as input for MARSHAL in order to calculate snapshots of the water uptake profile. In addition, RSHA generated by MARSHAL may be exported into formats readables by such models and therefore be readily used for such analysis.

The second main limitation of our analysis is that it does not account for the relative metabolic cost of the different metrics. For instance, one could argue that the carbon cost of producing more crown roots, both in terms of production and maintenance, is much larger than the cost of expressing more aquaporins (that could increase the root radial conductivity). Such estimations are estimated in some models at least for certain parameters (Postma et al. 2017) and could be included in a future version of MARSHAL. However, the inclusion of hydraulics in sensitivity analyses constitutes a step forward as compared to recent studies (Schnepf et al. 2018).

Furthermore, all the sensitivity analyses run in this study used uninformed uniform priors with an identical shape (boundaries given by the default parameters ± 50%). In the future, the model should take advantage of the existing literature to feed a bayesian meta-analysis in order to inform the actual parameter distributions and more accurately estimate the model uncertainty.

Finally, it must be kept in mind that the definition of an ideotype for drought tolerance should always be performed on a full plant model on specific environmental scenarios (Tardieu et al., 2017). Therefore, MARSHAL should still be coupled, for instance, to a shoot model, with atmospheric and soil boundary conditions representative of actual environments.

## Conclusion

MARSHAL is a novel numerical tool that couples different models and algorithms: a root architecture model (CRootBox), a solver of water flow in root system hydraulic architectures, a macroscopic property calculation method, and sensitivity and optimization functions. Its default input architectural and functional parameters were defined based on literature data but can be freely changed by users. Root system architectures can be exported in common format such as RSML by the model users. The generated root systems or their macroscopic properties can therefore be further used as inputs in other soil-root models, bridging the gap for virtual plant breeding.

Generated root system hydraulic architectures are compatible with macroscopic observations of the literature such as root length density profiles and root system conductance changes with time as demonstrated in two model validation examples in this study. This tool allows one bridging the gap between root local traits and plant upscaled hydraulic and architectural parameters.

Finally, sensitivity analysis highlighted sensitive local traits to macroscopic properties such as root system conductance (e.g. interbranching distances), depth of standard uptake (e.g. elongation rates), root system volume (root diameters) and root system convex hull (insertion angles) and allow reducing the dimensionality of optimization problem of root system hydraulic architecture in virtual root system breeding platforms.

Much attention was paid to making MARSHAL user friendly. As such, we created an R package (core function), and RMarkdown pipeline (batch analysis) and a web application (easy to use). MARSHAL code is released under an Open-Source licence.

## List of abbreviations

FSRSM: Functional Structural Root System Models
MARSHAL: MAize Root System Hydraulic Architecture soLver
RMSE: Root Mean Square of Error
RSHA: Root System Hydraulic Architecture
RWU: Root Water Uptake

## Model availability

MARSHAL is available as:

- an R package : https://github.com/MARSHAL-ROOT/marshal (https://doi.org/10.5281/zenodo.2391555)
- an online web application: https://plantmodelling.shinyapps.io/marshal/ (https://doi.org/10.5281/zenodo.2391249)
- a RMarkdown pipeline: https://github.com/MARSHAL-ROOT/marshal-pipeline (https://doi.org/10.5281/zenodo.2474420)

All relevant information can be found on the MARSHAL website: https://marshal-root.github.io/

